# Cytotoxicity and effector cooperation in the type III secretion system of *Aeromonas schubertii*

**DOI:** 10.1101/2025.03.21.644560

**Authors:** Hana Michova, Jan Pliva, Anezka Jirsova, David Jurnecka, Jana Kamanova

**Author notes:** Address correspondence to Jana Kamanova.

## Abstract

The type III secretion system (T3SS) is an important virulence factor of Gram-negative bacteria, including the genus *Aeromonas*, a group of aquatic bacteria capable of both mutualistic and pathogenic interactions. *Aeromonas* species are increasingly recognized as opportunistic human pathogens. The type strain *A. schubertii* ATCC 43700 encodes two distinct T3SSs located in the *Aeromonas* pathogenicity islands 1 and 2, hereby designated as API1 and API2, respectively. Presented work investigates the role of API1 and API2 in *A. schubertii*-induced cytotoxicity and identifies novel type III secretion effectors. HeLa cell infections showed that API1, but not API2, is essential for cellular cytotoxicity resulting in both apoptotic and necrotic cell death. The ΔAPI1 mutant failed to induce cytotoxicity, whereas the wild-type (WT) and ΔAPI2 strains induced comparable cytotoxic effects. Proteomic analysis identified 7 candidate effectors secreted by the API1 injectisome under low-calcium conditions. These included two previously characterized effectors, AopH and AopO of *A. salmonicida*, and five novel effectors hereby named AopI, AopJ, AopL, AopT, and AopU, whose injection into host cells via API1 was validated using a split luciferase reporter system. Functional analysis revealed distinct roles for these effectors. AopL, homologous to the VopQ effector of *Vibrio parahaemolyticus*, accelerated caspase 3-independent necrosis, while AopI, homologous to ExoY of *Pseudomonas aeruginosa*, suppressed caspase activation and necrosis, indicating a pro-survival function. These results show the role of API1 injectisome in the cytotoxicity of *A. schubertii* and expand our understanding of T3SS-mediated host-pathogen interactions in *Aeromonas* species.

**Importance:** This work demonstrates that the API1 injectisome is an important cytotoxicity determinant in *A. schubertii* and identifies novel effectors and their distinct contributions to host cell cytotoxicity, including the pro-survival effect of AopI and the cytotoxic effect of AopL. This interplay highlights a fine-tuned balance between pro-survival and cytotoxic mechanisms which is orchestrated by *A. schubertii* effectors.

## Introduction

The genus *Aeromonas* consists of heterogeneous group of aquatic bacteria that form long-term mutualistic relationships in the gastrointestinal tracts of fish and leeches (1–4). However, they are also prominent fish pathogens, causing substantial economic losses in aquaculture (5). In addition, *Aeromonas* species are increasingly recognized as opportunistic human pathogens, responsible for a wide range of diseases. Infections caused by *Aeromonas* range from mild gastroenteritis and wound infections to severe conditions such as necrotizing fasciitis and sepsis, particularly in immunocompromised individuals (6–10).

The genetic diversity and pathogenic potential of *Aeromonas* are closely linked to their ability to acquire mobile genetic elements, including plasmids, transposons and genomic islands that encode antibiotic resistance genes and virulence factors (11–13). Key virulence factors include structural components, such as flagella, pili and capsules, as well as a variety of secreted toxins and enzymes that disrupt host defenses. Among these, the type III secretion system (T3SS) and its effector proteins play a central role in the virulence of several *Aeromonas* species (14, 15).

The T3SS is a sophisticated nanomachine, often referred to as T3SS injectisome, that translocates effector proteins directly from the bacterial cytosol into host cells, where they manipulate host signaling pathways to benefit the bacteria. Structurally, the injectisome is composed of a basal body that spans the inner and outer membrane of the bacterium, an extracellular needle, and a translocon that forms a pore in the membrane of the host cell (16). The genes encoding these nanomachines are highly conserved, have evolved into seven families, and are located on mobile genetic element, termed pathogenicity island, which facilitate horizontal gene transfer (17–20). In contrast, genes encoding the effector proteins are more diverse and can be acquired independently, leading to their scattered distribution across the genome.

Previous studies have primarily focused on the Ysc family T3SS injectisomes and its effector proteins in *A. salmonicida,* as well as *A. hydrophila*, *A. piscicola* and *A. dhakensis* (20, 21). In *A. salmonicida*, the T3SS injectisome is encoded on a plasmid and its presence is essential for virulence of *A. salmonicida* in cold-water fish, with mutants showing reduced pathogenicity (22, 23). Injected effector proteins have diverse biochemical activities, including the bi-functional ADP ribosylating-GTPase activating effector AexT, the putative protein tyrosine phosphatase AopH, the phosphatidylinositolphosphatase Ati2, the putative acetyltransferase AopP and the putative serine/threonine kinase AopO, which were reported to affect cell viability, host immune responses, and cytoskeletal dynamics (24–29). Similarly, in *A. hydrophila, A. dhakensis* (formerly *A. hydrophila* SSU), and *A. piscicola* (formerly *A. hydrophila* AH-3), T3SS-positive strains exhibit enhanced virulence, while mutants show diminished cytotoxicity and increased susceptibility to phagocytosis. Insertional inactivation of T3SS injectisome leads to decreased cytotoxicity in carp epithelial cells, increased phagocytosis, and reduced virulence in blue gourami, rainbow trout or mice (30–33). However, only two effector proteins have been identified, AexT, which disrupts actin cytoskeleton and is homologous to *A. salmonicida*, and functionally similar AexU (34–37). Interestingly, a recent bioinformatic analysis of T3SS loci and effector proteins in the genomes of 105 *Aeromonas* strains revealed high variability and numerous potential effectors that are yet to be experimentally confirmed and further characterized. Moreover, some *Aeromonas* strains possess two distinct T3SS loci (38), hereby designated *Aeromonas* pathogenicity island 1 (API1) and *Aeromonas* pathogenicity island 2 (API2).

Presented work focuses on *A. schubertii*, which primarily affects aquatic animals. Infections with high mortality have been reported in commercially important species, such as whiteleg shrimp (39), tilapia (40), and asian seabass (41). Additionally, since 2009, a multi- drug resistant *A. schubertii* has emerged as a highly destructive pathogen in snakehead fish, *Channa maculata*, *C. argus* and their hybrid, which are widely cultured in southern China (42–44). The infection, known as “internal white spots disease”, causes white nodules scattered throughout the spleen, liver, and kidneys. These resemble mycobacterial granulomas and are likely formed as a result of *A. schubertii*-induced apoptosis and/or necrosis (45, 46). Outbreaks, which typically occur between May and October in intensive aquaculture settings, have resulted in severe economic losses in recent years (43, 44). In addition to its impact on aquatic species, *A. schubertii* is also an opportunistic pathogen of humans, associated with abscesses, wound infections, gastroenteritis and sepsis (47, 48). Intriguingly, *A. schubertii* lacks most of the T3SS effectors identified in other *Aeromonas* species, and the mechanisms responsible for its pathogenicity are poorly understood. Therefore, this study aimed to characterize the roles of API1 and API2-encoded injectisomes in *A. schubertii* infection using a HeLa cell model and identify effector proteins injected into host cells.

## Results

### *A. schubertii* carries two distinct T3SS injectisomes belonging to the Ysc and Ssa-Esc families

The type strain *A. schubertii* ATCC 43700 harbors two distinct T3SS loci, which are located on *Aeromonas* pathogenicity islands 1 (API1) and *Aeromonas* pathogenicity islands 2 (API2), and encode two separate injectisomes (Fig. 1A). Both loci contain all the structural proteins required for the assembly of a functional T3SS injectisome (Fig. 1B), however they differ in genetic organization and encoded proteins. While most of proteins encoded on API1 share high similarity with well characterized T3SS proteins from other bacteria, API2 harbors numerous hypothetical proteins with poorly characterized features (Fig. 1A). Additionally, similarity searches and phylogenetic analysis shown in Fig. 1C and Fig. S1 indicate that API1 belongs to the Ysc injectisome family, widely distributed in the *Aeromonas* genus, and closely resembling the injectisomes of *Yersinia* spp. and *Pseudomonas aeruginosa*. In contrast, API2 exhibits homology to the *Salmonella enterica* injectisome of the Ssa-Esc family, which is encoded on *Salmonella* pathogenicity island 2 (SPI2).

**Figure 1.**
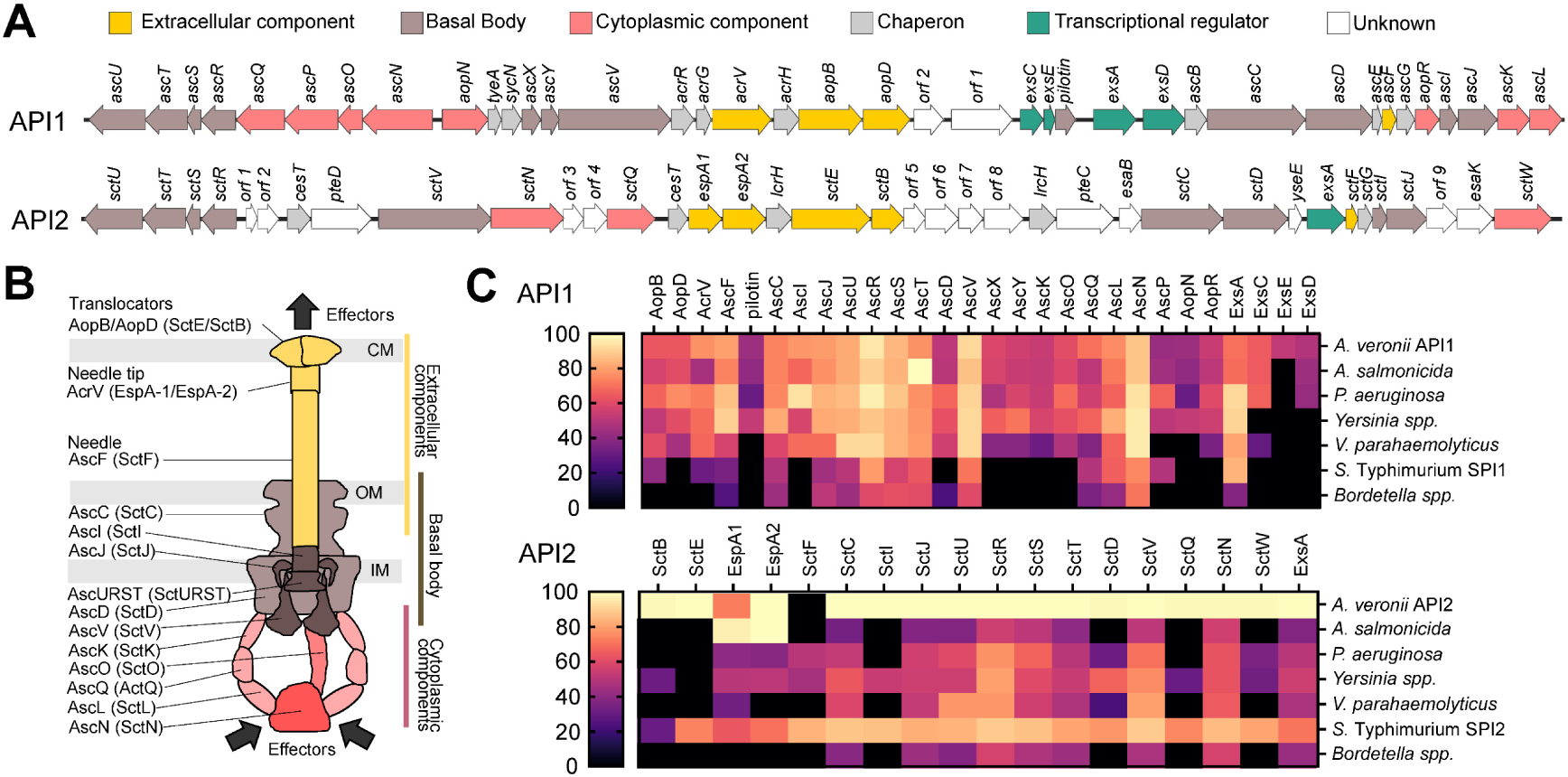
*Aeromonas schubertii* ATCC 43700 harbors two distinct loci encoding T3SS injectisomes. (A) Genetic organization of the API1 and API2 loci in *A. schubertii*. (B) Predicted structural organization of the T3SS injectisomes encoded by API1 and API2 loci. The diagram illustrates predicted localization of the proteins corresponding to the genes shown in panel (A). (C) Similarity of proteins encoded in the API1 and API2 loci to T3SS components from the indicated bacterial species. The heatmap displays the percentage similarity of each protein to its corresponding T3SS ortholog.

### API1 injectisome is required for cellular cytotoxicity of *A. schubertii*

To elucidate the role API1 and API2 injectisomes in the pathogenesis of *A. schubertii*, in-frame deletions of the ATPase genes *ascN* (API1) and *sctN* (API2) were generated, resulting in the mutant strains ΔAPI1 and ΔAPI2. These mutants were used together with the wild-type strain (WT) to infect HeLa cells with a multiplicity of infection (MOI) of 10:1. After one hour of infection, the extracellular bacteria were eliminated with gentamicin and time-lapse microscopy was performed. Morphological analysis revealed profound cellular changes and cytotoxicity of the WT strain, as shown in Fig. 2A and Video S1. Early changes included cell rounding, indicating modulation of the cytoskeleton and/or disruption of adhesion, followed by appearance of apoptotic features. These events culminated in cellular disintegration at later stages. Infection with the ΔAPI2 mutant led to similar effects as with the WT strain, while cells infected with the ΔAPI1 mutant did not show profound morphological changes and resembled uninfected controls (Fig. 2A and Video S1).

**Figure 2.**
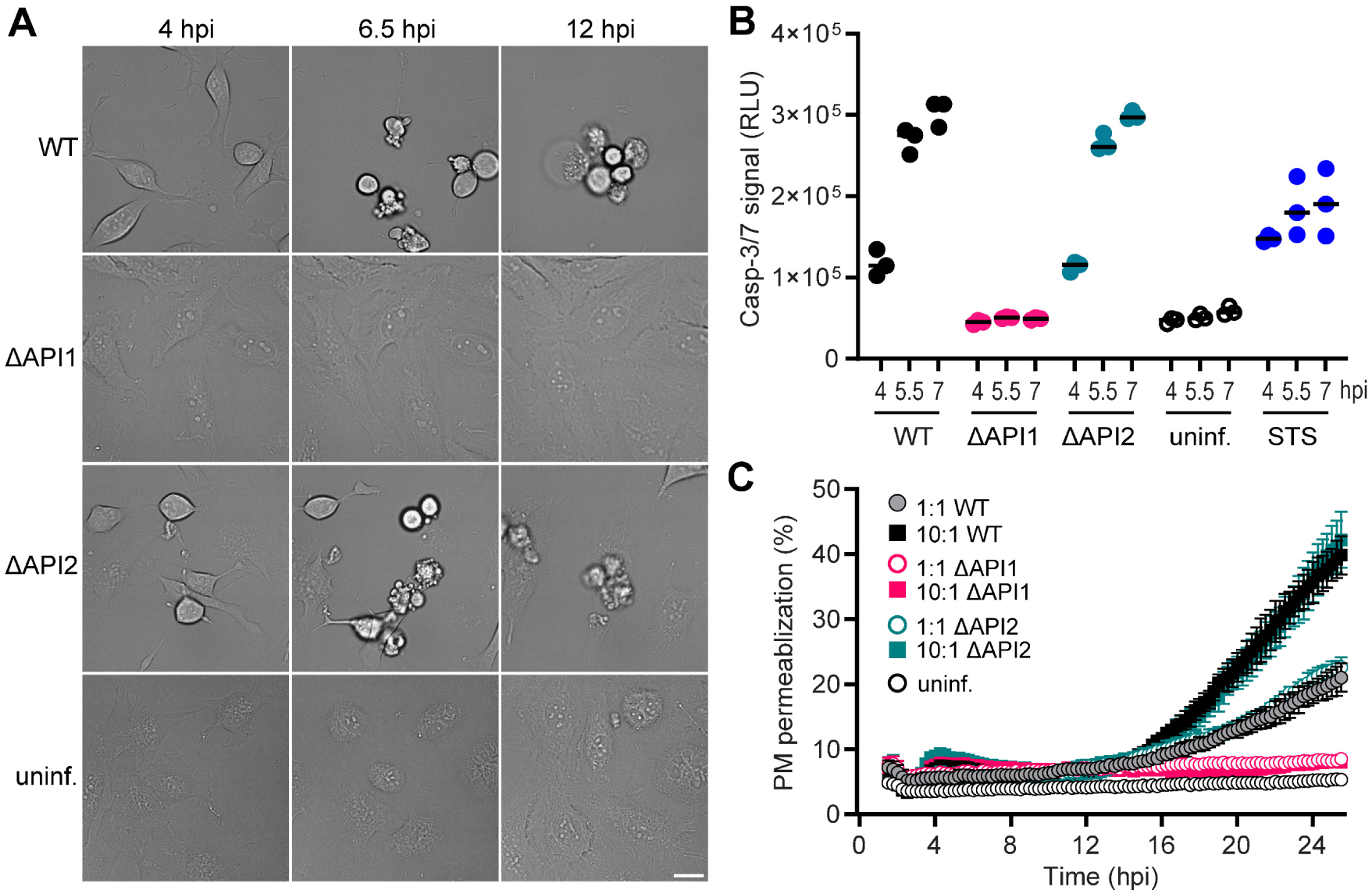
API1 injectisome is required for cellular cytotoxicity of *A. schubertii*. HeLa cells were either left uninfected or infected with *A. schubertii* wild-type (WT) or mutant strains lacking the API1 (ΔAPI1) or API2 (ΔAPI2) injectisomes, due to the deletion of the respective T3SS ATPases, at MOI 10:1. One hour post-infection, the extracellular bacteria were eliminated by addition of gentamicin. (A) HeLa cells were analyzed by live-cell imaging. A sequence of time lapse images is shown. Data are representative of 3 independent experiments. Scale bar, 20 µm. (B) Activation of caspase-3 and/or caspase-7 in infected HeLa cells was determined at indicated time points using the Caspase-Glo 3/7 assay, which detects cleavage of a proluminescent caspase-3/7 substrate. Data are presented as individual luminescence values (dots) from a representative experiment performed in triplicate wells out of 2 independent experiments. The black bar represents mean. HeLa cells treated with 1 µM staurosporine (STS) served as a positive control. (C) Real-time kinetics of plasma membrane (PM) permeabilization in infected HeLa cells was assessed using the fluorescent DNA-binding dye CellTox Green. PM permeabilization is expressed relative to complete permeabilization induced by a cell lysis solution. Data represent the mean ± SD of triplicate wells and are representative of 2 independent experiments.

To confirm these observations, we evaluated the induction of apoptosis by measuring caspase-3 and/or caspase-7 activity using the Caspase-Glo 3/7 assay, which quantifies luminescence generated by cleavage of a proluminescent substrate by active caspases. Besides, necrosis was assessed by examining plasma membrane permeabilization. As shown in Fig. 2B, infection with the WT and ΔAPI2 mutant strains at MOI of 10:1 resulted in caspase activation that was detectable as early as 4 hours post-infection, which was the earliest time point analyzed. Furthermore, the luminescence signal more than doubled within the following two hours, indicating increased caspase activity (Fig. 2B). In contrast, cells infected with the ΔAPI1 strain showed minimal caspase activity, comparable to uninfected controls. Western blot analysis confirmed these results and showed that caspase-3 was processed in HeLa cells infected with the WT and ΔAPI2 strains, but not in the cells infected with the ΔAPI1 strain (Fig. S2). In addition, plasma membrane permeabilization was observed in cells infected with the WT and ΔAPI2 strains at MOI of 10:1, as detected by the fluorescent DNA-binding dye CellTox Green, but only after 16 hours post-infection (Fig. 2C). Membrane disruption was also observed at a lower MOI of 1:1, albeit with lower efficiency. Both the WT and ΔAPI2 strains showed comparable necrosis, while cells infected with the ΔAPI1 mutant showed no signs of membrane permeabilization.

In summary, these results demonstrate that the API1 injectisome is essential for the delivery of effector proteins responsible for *A. schubertii*-induced cytotoxicity. This cytotoxicity includes activation of apoptotic caspases and cell necrosis. In contrast, the API2 injectisome does not contribute to cellular cytotoxicity in HeLa model system.

### Identification of candidate effectors of the API1 injectisome

To date, no T3SS effectors have been experimentally identified in *A. schubertii*. However, bioinformatic analysis of the *A. schubertii* CECT4240T genome uncovered homologs of two known *A. salmonicida* effectors, AopH and AopO, as well as several putative effectors (38). To identify API1-delivered effectors responsible for the cellular cytotoxicity, secretomes of WT and ΔAPI1 strains grown overnight in TSB medium were analyzed by mass spectrometry. However, no putative effectors were identified as significantly enriched in the secretome of WT strain at the set thresholds, suggesting that the API1 injectisome is inactive during standard cultivation condition (Fig. 3A). This aligns with the known activation mechanism of T3SSs, which are typically triggered upon contact with host cell membranes during infection. To overcome this limitation, TSB medium was supplemented with EGTA to chelate calcium ions and MgCl2 to repress PhoPQ system, which has been previously shown to artificially activate Ysc family of T3SS injectisomes (49–51). Under these conditions, API1 injectisome activity was robustly induced (Fig. 3B and Table S1). Comparative proteomic analysis identified numerous proteins secreted at significantly higher levels by the WT strain compared to the ΔAPI1 mutant. These included structural components of the API1 injectisome, homologs of the *A. salmonicida* effectors AopH (phosphotyrosine phosphatase) and AopO (serine/threonine kinase), and several candidate effectors including previously predicted putative effectors: PteI, hereby named AopI for *Aeromonas* outer protein I, homologous to the nucleotidyl cyclase ExoY of *Pseudomonas aeruginosa*; and PteJ, hereby named AopJ, homologous to the phosphothreonine lyase OspF of *Shigella flexneri* (38). Furthermore, two novel candidate effectors were identified and designated: AopT, containing a lipase domain, and AopU, harboring a RhoGAP domain (Table S2). Additionally, one more predicted potential effector PteL was detected in the WT strain, although its fold change was slightly below the selected fold change threshold (FC 3.552). As it is homologous to the VopQ effector of *Vibrio parahaemolyticus*, we decided to involve it in the further analyses and hereby named it as AopL.

**Figure 3.**
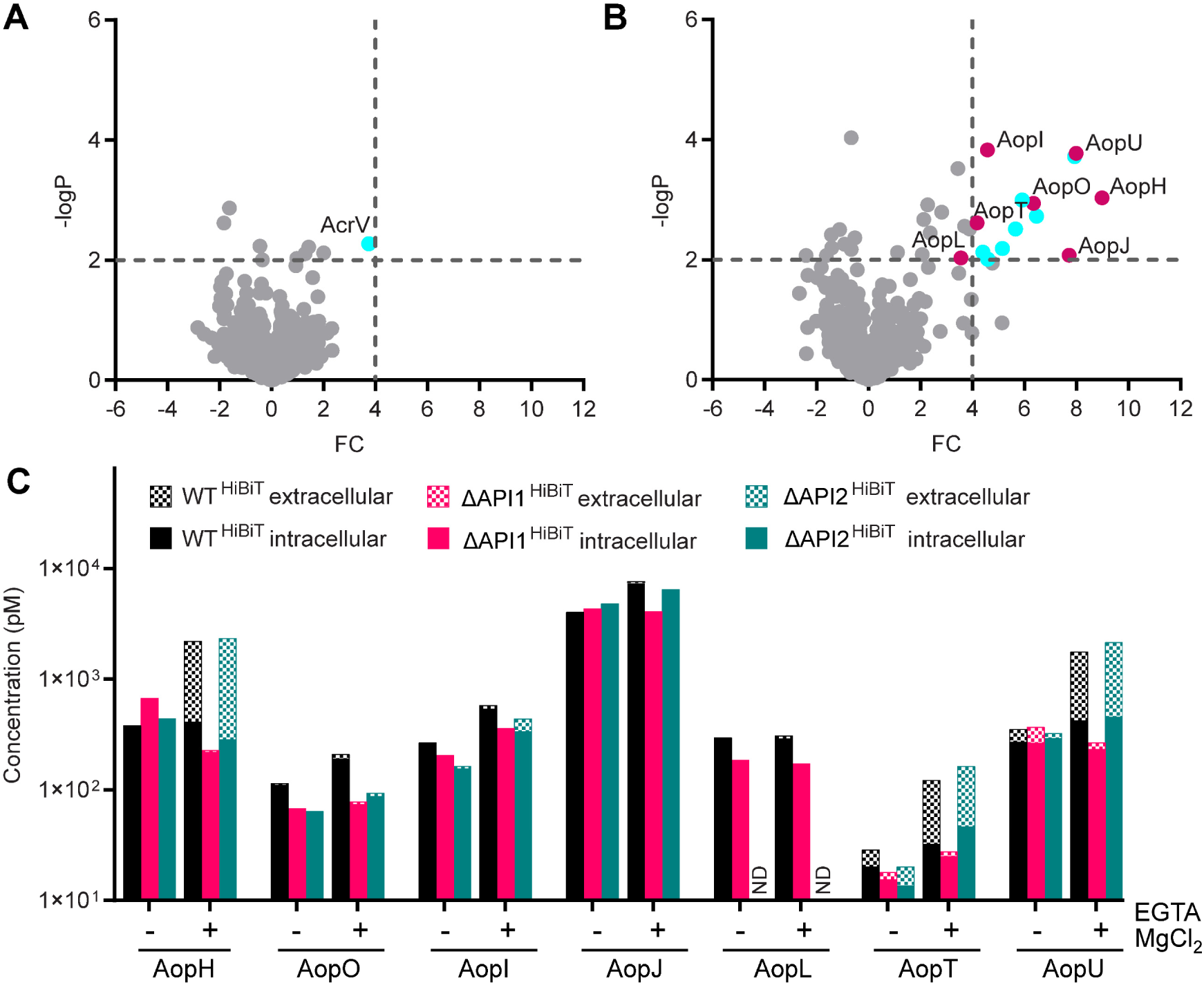
Identification of candidate effectors of the API1 injectisome. (A-B) Volcano plots illustrating global proteomic changes in the secretomes of *A. schubertii* wild-type (WT) compared to ΔAPI1 mutant strain lacking the API1 injectisome. Strains were grown in TSB medium without (A) and with supplementation (B) of 0.5 mM EGTA and 20 mM MgCl_2_. Red dots indicate candidate effector proteins, while blue dots represent proteins predicted to be structural components of the injectisome. Significant changes were defined as fold change ≥ 4 and -log_10_*P* value ≥ 2, corresponding to *P* ≤ 0.01. (C) Secretion of candidate effector proteins in response to low Ca^2+^/high Mg^2+^ concentrations. Luminescence measurements were used to quantify the extracellular and intracellular levels of candidate effector proteins. Measurements were performed on WT reporter (WT^HiBiT^) strains and the corresponding mutant derivatives lacking API1 (ΔAPI1^HiBiT^) and API2 (ΔAPI2^HiBiT^) injectisomes. Cultures were grown in TSB medium without and with supplementation of 0.5 mM EGTA and 20 mM MgCl_2._ Data represent the absolute concentration of candidate effector^HiBiT^ in extracellular and intracellular fractions and are the representative of 3 independent experiments. ND, not determined.

To validate the mass spectrometry results and quantify the presence of candidate effectors in both bacterial cells and secreted fractions, a split-luciferase system was employed. This approach utilizes high-affinity complementation between the 11-amino acid HiBiT tag on the C-terminus of the effector and an 18-kDa LgBiT fragment. Upon addition of the furimazine substrate, the reconstituted luciferase complex emits luminescence, allowing precise quantification of the tagged effector (52, 53). Reporter strains of *A. schubertii* were generated for each candidate effector by placing a HiBiT tag to the C-terminus of the effector in WT, ΔAPI1, and ΔAPI2 backgrounds, and luminescence was measured to assess candidate effector levels in intracellular and secreted fractions under normal and calcium-depleted / magnesium-high conditions. As shown in Fig. 3C (absolute candidate effector amounts) and Fig. S3 (extracellular fraction as a percentage of total), most candidate effectors were secreted through the API1 injectisome, after activation of secretion by calcium-depleted / magnesium- high conditions. Interestingly, AopH, AopT, and AopU displayed high secretion levels relative to their intracellular levels, indicating efficient export through API1 injectisome. In contrast, AopO, AopI, AopJ, and AopL showed minimal secretion, accumulating primarily intracellularly. Additionally, despite testing three independent clones, AopL^HiBiT^ expression was undetectable in the ΔAPI2 background for unknown reasons, preventing assessment of its secretion via API2.

In summary, secretion through API1 injectisome of *A. schubertii* is artificially activated in *vitro* under low-calcium conditions coupled with high magnesium concentration. Additionally, we experimentally identified several predicted and two novel candidate effectors in *A. schubertii* ATCC 43700, which are secreted through API1 injectisome.

### AopI, AopJ, AopL, AopT, and AopU represent novel effectors of *Aeromonas* species

To further corroborate the *in vitro* secretion data, we tested whether the candidate effector proteins were injected into the host cells. The HiBiT-tagged effector reporter strains were used to infect HeLa cells expressing LgBiT at MOI of 50:1 in the presence of a cell-permeable furimazine substrate. Successful translocation was indicated by an increase in luminescence (translocation signal) emitted due to the complementation of HiBiT tag and LgBiT subunit constitutively expressed in the host cell cytosol (52, 53).

As shown in Fig. 4A, luminescence was detected within the first hour of infection for all seven candidate effectors of *A. schubertii* ATCC 43700, demonstrating their injection into the host cytosol. These included the previously identified *A. salmonicida* effectors AopH and AopO, as well as the newly identified effectors AopI, AopJ, AopL, AopT, and AopU. Effector delivery was exclusively mediated by the API1 injectisome, except for AopL, which could not be tested due to undetectable AopL^HiBiT^ expression in the ΔAPI2 background (see above). Interestingly, the amount of translocation signal for respective effectors (Fig. 4B) did not always correlate with their *in vitro* secretion levels. While AopH and AopU were both efficiently secreted and translocated, AopO exhibited and unexpectedly high translocation signal despite its lower secretion levels in *A. schubertii* culture. In contrast, AopT, which was efficiently secreted in *vitro*, showed weak translocation signal. The remaining effectors, AopJ, AopI, and AopL, displayed translocation signals consistent with their *in vitro* secretion levels.

**Figure 4.**
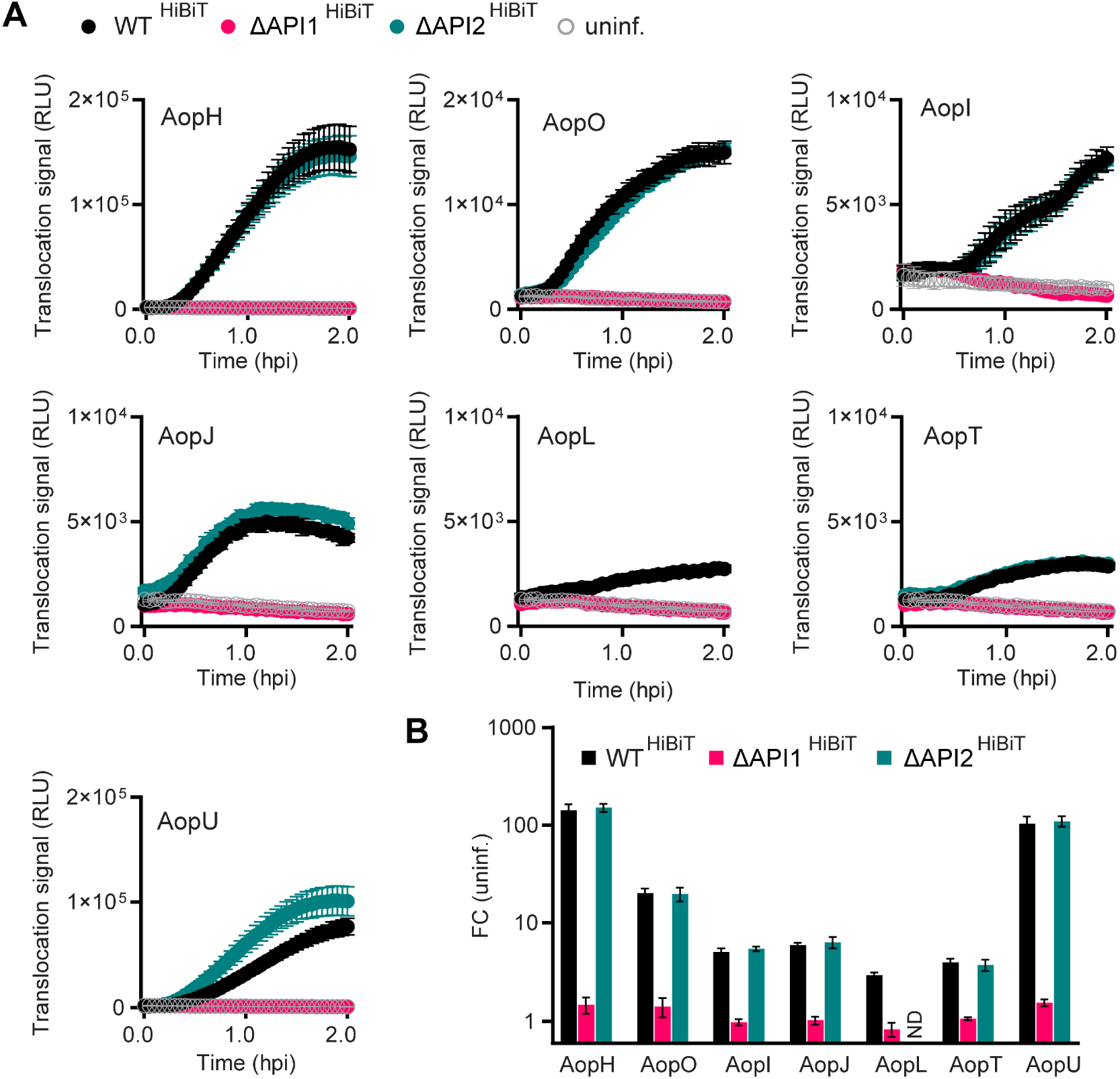
AopI, AopJ, AopL, AopT and AopU represent novel effectors of *Aeromonas* species. LgBit-expressing HeLa cells were infected with wild-type reporter strains (WT^HiBiT^) or mutant reporter strain derivatives lacking API1 (ΔAPI1^HiBiT^) or API2 (ΔAPI2^HiBiT^) injectisomes at MOI 50:1. Infections were performed in the presence of a cell-permeable furimazine substrate. Luminescence was measured every 3 min and expressed as relative luminescence units (RLU). (A) Data represent the mean ± SD of triplicate wells from a representative experiment out of three independent experiments. The candidate effector^HiBiT^ is indicated. ND, not determined for *aopL*^HiBiT^ /ΔAPI2. (B) Fold change of luminescence for cells infected with reporter strains relative to uninfected cells. Data represent the mean fold change in maximum luminescence ± SD, calculated from three independent experiments, each performed in triplicate.

Overall, these data demonstrate that AopI, AopJ, AopL, AopT and AopU are novel effectors within *Aeromonas* genus.

### Pro-survival effector AopI counteracts the cytotoxic effects of AopL in HeLa cells

To investigate the role of the newly identified effectors in host cell cytotoxicity, individual effector genes were deleted from the genome of *A. schubertii*, and the resulting mutant strains were tested for their effects on caspase activation and plasma membrane integrity. Remarkably, most of the mutant strains were comparable to WT, both in caspase-3/7 activation at 7 hours post-infection and overall disruption of the plasma membrane integrity (Fig. 5A-D). Even the double mutant Δ*aopH*/Δ*aopU*, which lacks the genes encoding the highly translocated effectors, did not significantly alter overall cytotoxicity (Fig. 5C). However, Δ*aopI* mutant showed a significant increase in caspase activity (Fig. 5A), which correlated with enhanced cellular necrosis at later stages of infection as determined by the CellTox assay (Fig. 5B). This indicates that AopI suppresses caspase activation during infection and postpones the overall cytotoxic effect of API1. Moreover, Δ*aopL* mutant exhibited a delay in plasma membrane permeabilization, which occurred approximately 5 hours later than in the WT strain (Fig. 5D). This delay was not accompanied by differences in caspase 3/7 activation, suggesting that AopL facilitates a caspase-3/7-independent necrotic pathway. In contrast, the Δ*aopU* mutant showed a slight but significant decrease in caspase-3/7 activation, which, however, did not translate into changes in the overall cellular necrosis (Fig. 5A and 5C).

**Figure 5.**
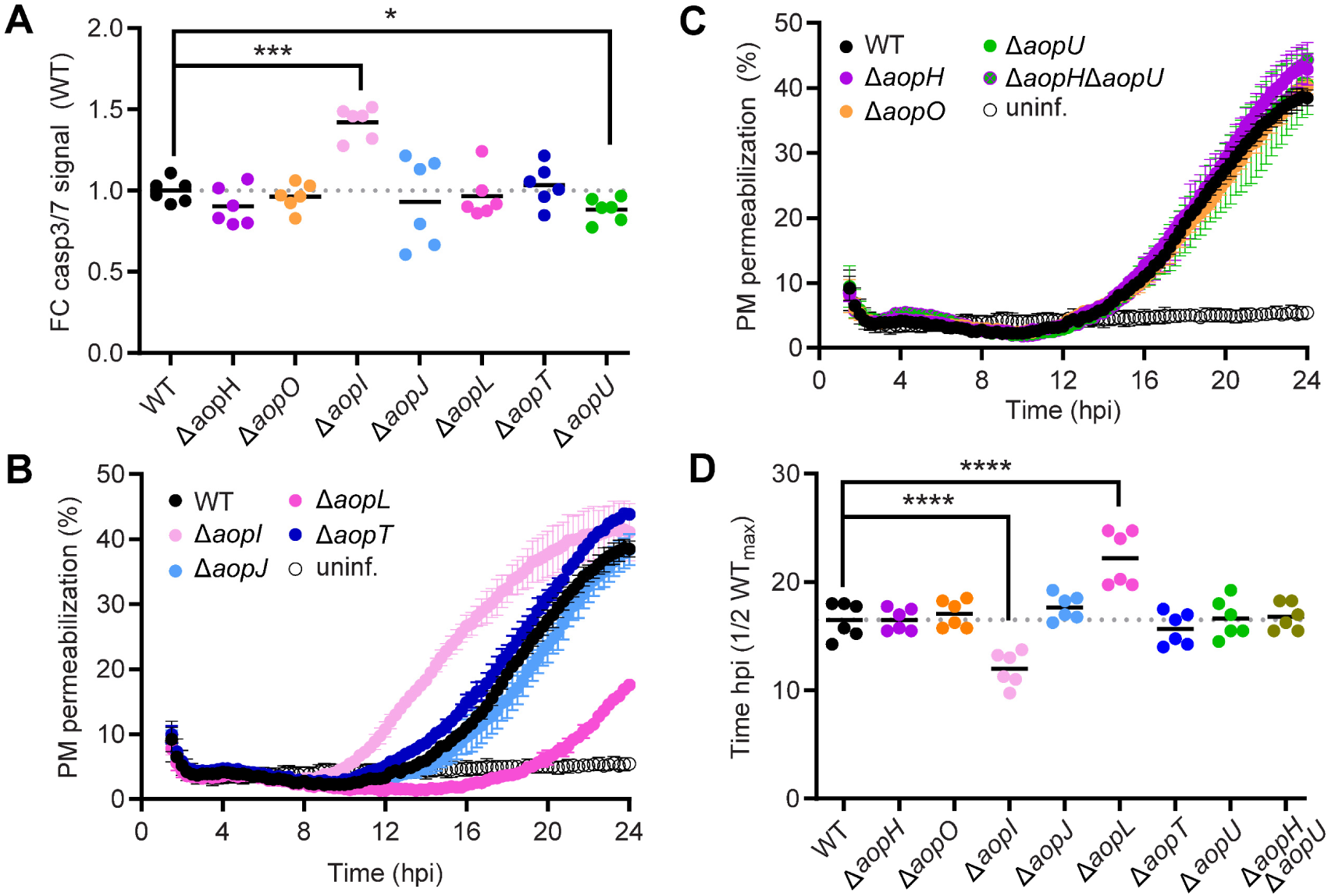
Pro-survival effector AopI counteracts the cytotoxic effects of AopL in HeLa cells. HeLa cells were either left uninfected or infected with *A. schubertii* wild-type (WT) or mutant strains lacking specific API1 effectors (Δ*aopH*, Δ*aopO*, Δ*aopI*, Δ*aopJ*, Δ*aopL*, Δ*aopT* and Δ*aopU)* or their combination (Δ*aopH* Δ*aopU*) at MOI 10:1, as indicated. One hour post-infection, the extracellular bacteria were eliminated by addition of gentamicin. (A) Caspase-3 and/or caspase-7 activation in infected HeLa cells was measured at 7 h post-infection using the Caspase-Glo 3/7 assay, which detects cleavage of a proluminescent caspase-3/7 substrate. Data are presented as the fold change (FC) of luminescence of cells infected with mutant strains relative to those infected with wild-type strain. Data represent two independent experiments, each with triplicate wells. \**p* < 0.05, \*\*\**p* < 0.001, unpaired two-tailed *t-*test. (B-C) Real-time kinetics of plasma membrane (PM) permeabilization of infected HeLa cells was measured using the fluorescent DNA-binding dye CellTox Green. PM permeabilization is expressed relative to complete permeabilization induced by a cell lysis solution. Data represent the mean ± SD of triplicate wells and are representative of 2 independent experiments. (D) Statistical analysis of PM permeabilization. Data are expressed as the time required to reach 50% of the maximal permeabilization observed in the WT-infected cells. Values represent two independent experiments, each performed in triplicate wells. ****p < 0.0001, unpaired two-tailed *t*-test.

Altogether, these results emphasize the critical role of AopL in accelerating cell death, albeit via a mechanism that primarily does not involve caspase-3/7 activation. In contrast, AopI acts as a pro-survival effector that counteracts the induction of cell death pathways during infection.

## Discussion

This study provides new insights into the role of the T3SS and its effector proteins in *A. schubertii* pathogenesis. Using a HeLa cell model, we demonstrated that the API1 is required for *A. schubertii*-induced cytotoxicity, whereas API2 plays a negligible role in the process. This distinction is consistent with previous observations that bacterial genomes often harbor multiple T3SS loci, each with specialized functions that depend on the environmental context.

The API1, which encodes API1 injectisome of *A. schubertii,* belongs to the Ysc family of injectisomes that also includes injectisomes of *Yersinia* spp., *P. aeruginosa* and *Vibrio* spp. (20). Our findings indicate that it can be activated by artificial cues, particularly low Ca^2+^ and high Mg^2+^ concentrations, likely through both transcriptional and post-translational regulation. Similar to *P. aeruginosa*, API1 encodes components of the ExsACDE regulatory circuit. In *P. aeruginosa*, T3SS expression is controlled by ExsA, a transcriptional activator which is bound to a negative regulator ExsD under non-permissive conditions. Upon exposure to inducing signals, such as host cell contact or low Ca^2+^ level, ExsC sequesters ExsD, thereby freeing ExsA to activate T3SS gene expression (54, 55). Additionally, low Ca^2+^ serves as an artificial activation signal by triggering the release of the gatekeeper protein from the SctV component in various injectisomes, including those of *Yersinia*, and *Bordetella,* which subsequently allows for effector secretion in the so-called second substrate switch event (50, 51, 53, 56). High Mg^2+^ concentrations, on the other hand, have been reported to repress the two-component PhoPQ system, thereby upregulating transcription of API1 machinery (49). However, the exact roles of low Ca^2+^ and high Mg^2+^ concentration in API1 activation of *A. schubertii* remain to be fully established.

By depleting Ca^2+^ and increasing Mg^2+^ to artificially activate the API1 injectisome, we identified seven candidate effectors through mass spectrometry. Among these, two (AopH and AopO) are homologous to effector proteins in *A. salmonicida* (29), three (AopI, AopJ, and AopL) were predicted in a recent bioinformatic analysis (38), and two (AopT and AopU) are novel. Interestingly, our mass spectrometry analysis did not detect all bioinformatically predicted API1 T3SS effectors (38), including PteA, which is homologous to cytotoxic effector BteA of *Bordetella* species (57, 58). Indeed, *pteA* gene is not pseudogenized in *A. schubertii* ATCC 43700, and a T3SS chaperone is located upstream of its coding region, supporting its classification as a T3SS effector. This suggests that alternative induction conditions, such as growth at 37°C or deletion of the ExsD negative regulator, may reveal additional T3SS effectors of API1 injectisome.

To validate translocation of the identified candidate effector into host cells, we employed a split-luciferase system (52, 53). Intriguingly, translocation signals did not always correlate with intracellular abundance or secretion levels observed *in vitro*. For instance, AopO exhibited an unexpectedly high translocation signal, suggesting the involvement of additional regulatory factors mediating its preferential delivery, or increasing expression after direct contact with a host cell. Alternatively, variations in HiBiT-LgBit complementation efficiency in bacterial supernatants vs. host cell cytosol, potentially influenced by the accessibility of the HiBiT-tag to LgBit subunit, may have also contributed.

Although the observed cell cytotoxicity was API1-dependent, absence of individual API1 effectors did not entirely prevent cell death. This suggests a degree of redundancy or functional overlap among the API1 effectors, which is quite common for T3SS effectors (59). Among the 7 identified effectors, only AopL and AopI were key modulators of HeLa cell death, though in contradictory manner, which indicates a functional dichotomy. Specifically, AopL accelerated necrosis through a caspase-3-independent mechanism, which resembled the effect of its homolog, VopQ in *Vibrio parahaemolyticus*. This effector targets lysosomal V-ATPase of host cell and triggers multiple detrimental effects, including lysosomal deacidification, disruption of redox homeostasis, and induction of autophagy and cell necrosis (60–63). In contrast, AopI exhibited a protective, pro-survival effect by suppressing caspase-3/7 activation. This protein is homologous to ExoY of *P. aeruginosa*, which is a nucleotidyl cyclase that is activated by filamentous actin (64). Interestingly, ExoY has been shown to promote virulence by modulating innate immune responses as well as to protect against *P. aeruginosa*- induced cytotoxicity in human bronchial epithelial cells (65, 66).

Next to these key modulators, AopU also played a minor yet significant role in caspase- 3/7 activation, although its precise function remains unclear. AopU is predicted to inactivate Rho, Rac and/or Cdc42 through GTP hydrolysis mediated by its putative RhoGAP activity, similar to *Aeromonas* AexT/AexU (37, 67). This predicted activity aligns with the early morphological changes observed during *A. schubertii* infection, which were characterized by cell rounding, likely resulting from disruption of cell cytoskeleton and/or loss of adhesion. However, unlike AexT/AexU, AopU lacks ADP-ribosyltransferase domain. Additionally, AopH, a putative phosphotyrosine phosphatase, and AopO, a putative serine/threonine kinase, may also contribute to these morphological changes. Their *Yersinia* homologs, YopH and YopO, are known to alter actin cytoskeleton, inhibit phagocytosis and disrupt focal adhesions (29).

In addition, among the newly identified effectors, AopJ might contribute to disrupting innate immune signaling, as it is homologous to a family of phosphothreonine lyase effectors, including *Shigella* OspF and *Salmonella* SpvC, which inhibit MAPK signaling pathways and downregulate inflammatory cytokine production (68, 69). Meanwhile, the probable function of AopT, a putative lipase, remains unknown. Future studies should investigate how the identified effectors influence macrophage phagocytic activity and susceptibility to predation by amoebas, as well as their potential roles in subversion of innate or adaptive immune responses in fish and humans. Given the increasing threat of *A. schubertii* to aquaculture of snakehead fish, it will be also critical to assess whether the T3SS and its effectors contribute to the formation of white nodules in the spleen, liver, and kidneys of diseased animals, and overall fish morbidity (45, 46, 70).

All identified T3SS effectors, except for AopL, which could not be tested, were translocated into the host cell specifically through API1 injectisome. This specificity may result from the intrinsic selectivity of the API1 secretion system, or the inactivity of the API2 injectisome during HeLa cell infection. API2 shares homology with the *Salmonella enterica* injectisome of the Ssa-Esc family, which is activated after bacterial uptake into acidic compartments by low pH (71). Although *A. schubertii* did not appear to actively invade HeLa cells (data not shown), and is generally considered an extracellular pathogen, other *Aeromonas* species have been reported to invade epithelial cells and amoebas (72–75). Therefore, it remains important to determine whether the API2 injectisome functions extracellularly or is activated only upon bacterial uptake, potentially facilitating intracellular persistence.

Overall, our results emphasize the similarity between the API1 effectors of *A. schubertii* and T3SS effectors of other Gram-negative bacteria, in particular *A. salmonicida*, *P. aeruginosa* and *V. parahaemolyticus*, which supports the idea of horizontal gene transfer within *Aeromonas* genus (11–13). However, it is critical to mention that the identified set of API1 effectors is unique for *A. schubertii*. This suggest that *A. schubertii* may have acquired two distinct APIs and their effectors through horizontal gene transfer, and retained them for a competitive advantage in a specific ecological niche. As such, our study advances understanding of *A. schubertii*-mediated cytotoxicity in the HeLa cell model and provides experimental identification of novel *Aeromonas* effectors.

## Material and methods

### Bacterial strains and growth conditions

The type strain of *Aeromonas schubertii* ATCC 43700 (CDC 2446-81T) originally isolated from a human forehead abscess in Texas (76), together with its derived mutant strains, were used throughout this study. A detailed list of strains is provided in Table S3. Bacteria were cultivated on tryptone soya agar (TSA; Oxoid) at 30 °C for 48 h following inoculation from stocks preserved in 40% glycerol at -80 °C. Liquid cultures of *A. schubertii* strains were grown in tryptone soya broth (TSB; Oxoid) at 30 °C with constant shaking at 180 rpm. For experiments, overnight cultures in TSB were subcultured at a 1:25 ratio into fresh TSB and incubated for approximately 4 h to reach the exponential growth phase unless stated otherwise. For plasmid construction, *Escherichia coli* strain XL-1 Blue was used, while *E. coli* strain SM10λ pir was employed for plasmid transfer into *A. schubertii* via bacterial conjugation. *E. coli* strains carrying the temperature-sensitive allelic exchange vector pAX2 were cultured at 30 °C on LB agar or in LB broth. When appropriate, LB media were supplemented with 100 µg/mL ampicillin.

### Plasmid construction and generation of *A. schubertii* mutant strains

Mutant strains of *A. schubertii* ATCC 43700, including in-frame deletion mutants and HiBiT-tagged strains, were generated through homologous recombination using the pAX2 allelic exchange vector (gift from Karen Guillemin; Addgene plasmid # 117398; http://n2t.net/addgene:117398; RRID: Addgene_117398; (77)). Details of pAX2-derived plasmids used in this study are listed in Table S4. Plasmids were constructed using the Gibson assembly method (78), in which PCR-amplified DNA fragments derived from chromosomal DNA of *A. schubertii* ATCC 43700 were ligated into the *Sma*I-linearized pAX2 vector.

Constructed plasmids were verified by DNA sequencing (Eurofins Genomics) before being introduced into *A. schubertii* by bacterial conjugation. For conjugation, *A. schubertii* and *E. coli* SM10λ pir strain were mixed in a 1:1 ratio on a filter disk and placed on TSA. The mating mixture was incubated overnight at 30 °C. After the incubation, bacteria were recovered and plated on TSA supplemented by gentamicin (10 µg/ml) and anhydrotetracycline (10 ng/ml) to select for *A. schubertii* merodiploids, as described previously (77). Isolated merodiploid colonies were screened for the loss of GFP expression indicating the second recombination event. PCR genotyping was performed to differentiate between wild-type and mutant alleles.

### Mammalian cell culture

Cell lines HeLa (ATCC CCL-2, human cervical adenocarcinoma) and HeLa-LgBit (HeLa cells constitutively expressing LgBit; (53)) were cultivated in Dulbecco’s Modified Eagle Medium supplemented with 10% (vol/vol) heat-inactivated fetal bovine serum (DMEM-10% FBS) at 37 °C and 5% CO_2_.

### Time-lapse HeLa live cell imaging

To visualize host cells during *A. schubertii* infection, 1 x 10^5^ of HeLa cells in DMEM-10% FBS were seeded to each well of 4-well glass-bottom dish (Cellvis) and allowed to adhere overnight. The next day, cells were either left uninfected, infected with *A. schubertii* wild type (WT) or its ΔAPI1 and ΔAPI2 derivatives (Table S3) at MOI of 10:1. To enhance infection efficiency, the dish was briefly centrifugated (5 min; 120 g). One hour post-infection, gentamicin was added to a final concentration of 100 µg/ml to stop the infection, and dish was transferred to a prewarmed stage-top incubation chamber of motorized fluorescence microscope (IX-83, Olympus, Japan) equipped with sCMOS camera Photometrics Prime 95B. Bright-field images were acquired using a 40x dry objective (UPLXAPO40X, NA = 0.95) under controlled conditions of 37 °C and 5% CO2. Sequential 16-bit images were acquired as a time-lapse of 24 hours with frame intervals of 10 minutes using the CellSens software. Images were then processed with ImageJ (Fiji, (79)) by enhancing the brightness to contrast ratio, selecting ROI and adding scale bar.

### Determination of caspase 3 and/ or caspase 7 activation using Caspase-Glo assay

To evaluate caspase-3 and/or -7 activation during infection, HeLa cells (1.5 x 10^4^ per well) were seeded in a 96-well plate in DMEM-10% FBS. The next day, *A. schubertii* strains grown to exponential phase were washed twice with DMEM-10% FBS by centrifugation (3 min; 8,000 g) and added to cells at MOI 10:1. The plate was centrifuged (5 min; 120 g) to enhance the infection efficiency. One hour post-infection, infection was stopped by addition of gentamicin (100 µg/ml), and cells were further incubated at 37 °C with 5% CO2. At designated time points, an equal volume of freshly reconstituted Caspase-Glo 3/7 Reagent (Promega, Cat. No. G8091), containing a pro-luminescent caspase-3/7 substrate, was added to each well. The plate was then incubated for 30 min at 37 °C to allow cell lysis and substrate cleavage by activated caspases, generating luminescent signal. Following the incubation, 100 µL of the reaction mixture from each well was transferred to a 96-well white/clear bottom plate (Corning). Luminescence was measured using The Spark microplate reader (Tecan) with an integration time of 1 s per well. Experimental conditions included both uninfected cells, and a positive control treated with 1 µM staurosporine. For statistical analysis, data were normalized by the average luminescence of WT-infected samples and compared using two-tailed unpaired parametric t-test.

### Analysis of caspase 3 activation by Western Blot

HeLa cell were seeded in 6-well cell culture plate (5 x 10^5^ per well), and infected the following day with *A. schubertii* strains washed in DMEM-10%FBS at MOI 10:1. The plate was centrifugated (5 min; 120 g), and infection was terminated after 1 h by addition of gentamicin (100 µg/ml). At indicated time points, the content of each well was aspired, and cells were then lysed by addition of 100 µl of lysis buffer consisting of 0.3% (vol/vol) Triton X-100 in PBS containing the Complete Protease Inhibitor Cocktail (Roche). Extracted proteins were boiled (95 °C; 5 min), separated by 15% SDS-PAGE electrophoresis and transferred onto a nitrocellulose membrane. Membranes were probed overnight with rabbit polyclonal antibody against caspase-3 (dilution 1:1,000; CST, Cat. No. 9662). The detected caspase-3 was revealed with 1:3,000-diluted horseradish peroxidase (HRP)-conjugated anti-rabbit IgG secondary antibodies (GE Healthcare) using a Pierce ECL chemiluminescence substrate (Thermo Fisher Scientific) and Image Quant LAS 4000 station (GE Healthcare). For loading control, membranes were re-probed with mouse monoclonal IgG against β-actin (dilution 1:100,000; Proteintech, Cat. No. 66009-1-Ig) and revealed with secondary HRP-anti-mouse IgG (Cytiva), as above.

### Determination of plasma membrane permeabilization using CellTox Green assay

HeLa cells (2 x 10^4^ per well) were seeded in a 96-well black/clear bottom plate (Corning) in DMEM-10% FBS. The next day, *A. schubertii* and the derived mutant strains were washed in DMEM-10% FBS and added to the wells at the indicated MOI. The plate was then centrifugated (5 min; 120 g) to facilitate bacterial contact with cells. One hour post-infection, gentamicin (100 µg/ml) and the fluorescent DNA-binding dye CellTox Green (Cat. No. G8743, Promega) were added to each well. The plate was subsequently incubated under controlled conditions (37°C and 5% CO₂) inside The Spark microplate reader (Tecan). Fluorescence measurements were recorded at 15 min intervals over a 24 h using excitation and emission wavelengths of 490 nm and 525 nm, respectively. For statistical analysis, the time at which the WT signal reached half of its maximum was compared to the corresponding time for mutant strains using a two-tailed unpaired parametric *t-*test.

### Proteomic analysis of secretomes

Bacterial cultures were grown overnight in 50 ml of TSB medium with or without supplementation of 0.5 mM EGTA and 20 mM MgCl2 at 30 °C with constant shaking of 180 rpm. Subsequently, cells were removed by centrifugation (30 min; 10,000 g; 4 °C) followed by filtration through 0.22 μm membranes. The resulting supernatants containing secreted proteins were precipitated by adding trichloracetic acid (Sigma) to final concentration of 10% (v/v) and incubating at 4 °C overnight. Precipitated proteins were collected by centrifugation (30 min; 14,000 g; 4 °C), washed twice with cold acetone, air-dried and dissolved in 50 mM ammonium bicarbonate with 8M urea. Protein concentrations were determined using the Pierce BCA Protein Assay Kit (Thermo Fisher Scientific), and 70 µg of protein per sample was used for further processing. Samples were analyzed by label-free mass spectrometry (MS) using a nanoLC-MS/MS system coupled to an Orbitrap Fusion Tribrid mass spectrometer (Thermo Fisher Scientific) as previously described (80). Data were analyzed and quantified using a label-free quantification approach with MaxQuant (version 2.5.2.0) and Perseus (version 2.0.11). The false discovery rate (FDR) was set to 1% for both proteins and peptides. The enzyme specificity of trypsin was set as C-terminal to Arg and Lys residues. Carbamidomethylation was set as a fixed modification, while N-terminal protein acetylation and methionine oxidation were considered variable modifications. The maximum number of missed cleavages was set to two. Protein identification was performed using the *Aeromonas schubertii* reference proteome database (*Aeromonas schubertii* ATCC 43700, Uniprot - UP000054876). Statistical analysis was conducted in Perseus using a student t-test with a significance threshold of p < 0.01, and only proteins with a fold change ≥ 4 were considered upregulated. The mass spectrometry proteomics data have been deposited to the ProteomeXchange Consortium via the PRIDE (81) partner repository with the dataset identifier PXD062075.

### Quantification of intracellular and secreted levels of candidate effectors by luminescence measurements

To evaluate the levels of secreted and intracellular effectors, reporter strains expressing HiBit-tagged candidate effectors and their ΔAPI1 and ΔAPI2 derivatives (Table S3) were cultivated in TSB medium with or without supplementation of 0.5 mM EGTA and 20 mM MgCl_2_ to reach OD600nm 1 ± 0.05. One ml of cultures was centrifuged (5 min; 8,000 g) and levels of candidate effectors were quantified in both the supernatants representing the secreted fraction, and in the cell pellet extracts containing intracellular effectors. The quantification was performed using the Nano-Glo HiBiT Extracellular Detection System (Promega, Cat. No. N2420) as previously described (53).

In brief, for preparation of cell extracts, cell pellets were resuspended in 200 µl of extraction buffer (150 mM NaCl in 50 mM Tris-HCl, pH 8.0) and combined with 0.1 mm glass beads (Scientific Industries). Cells were lysed using the Disruptor Genie (Scientific Industries) with two cycles of bead beating (3 min at maximum speed) followed by cooling interval of 3 min on ice. The disrupted cell suspension was diluted with 800 µl of the same buffer and clarified by centrifugation (10 min; 14,000 g). Cell culture supernatants and pellet extracts were subsequently transferred into 96-well white/clear bottom plate and mixed with an equal amount of reconstituted Nano-Glo HiBiT Extracellular Reagent containing recombinant LgBit protein and furimazine substrate in Nano-Glo buffer. Luminescence was measured using The Spark microplate reader (Tecan) with an integration time of 1 s per well. To determine the exact concentration of effectors, serial dilutions of purified recombinant LRT-HiBiT protein were included in each plate as a standard (53).

### Determination of effector injection into the host cell

To evaluate the injection of candidate effectors into the host cells, 2x 10^4^ of HeLa-LgBit (HeLa cells constitutively expressing LgBit) per well were seeded in 96-well white/clear bottom plate (Corning), in DMEM-10% FBS. The next day, reporter strains expressing HiBit-tagged candidate effectors and their ΔAPI1 and ΔAPI2 derivatives (Table S3) grown to exponential phase were washed twice in DMEM-10%FBS by centrifugation (5 min; 8,000 g) and added to cells at MOI of 50:1, along with Nano-Glo Live Cell Reagent (Promega, Cat. No. N2011) containing cell permeable luciferase substrate in Nano-Glo buffer. After centrifugation (5 min; 120 g), the plate was placed inside the chamber of The Spark microplate reader (Tecan) with 37 °C and 5% CO2 and luminescence measurements were performed for 2 h at 3 min intervals with integration time 1 s. The maxima of the signal reached for each tagged effector were normalized as a fold change towards uninfected control and were statistically evaluated using two-tailed unpaired parametric t-test.

### Bioinformatic analysis

The genome sequence of *A. schubertii* ATCC 43700 was downloaded from NCBI (BioProject PRJNA304368). Homology of T3SS-related proteins and potential effectors was established using Protein BLAST run against non-redundant protein sequences of *Aeromonas salmonicida*, *Aeromonas veronii*, *Yersinia* spp., *Vibrio parahaemolyticus*, *Pseudomonas aeruginosa*, *Salmonella enterica*, and *Bordetella* spp. Only proteins with the highest Percent Identity were taken into consideration.

To create a phylogenetic tree, sequences of SctN and SctC from various T3SS-expressing bacteria were downloaded from Uniprot database (Table S5). Sequences were aligned and their phylogenetic tree was constructed as Minimum Evolution Tree based on p-distance using MEGA 11 software (82). Trees were further annotated using online tool iTOL v7 (83).

## Acknowledgements

This work was supported by grant Talking microbes - understanding microbial interactions within One Health framework (CZ.02.01.01/00/22_008/0004597) of the Ministry of Education, Youth and Sports of the Czech Republic (www.msmt.cz) and the Lumina Queruntur Fellowship LQ200202001 of the Czech Academy of Sciences to J.K. We also acknowledge the support of the project LM2023053 (Czech National Node to the European Infrastructure for Translational Medicine) from the Ministry of Education, Youth and Sports of the Czech Republic (www.msmt.cz). The funders had no role in study design, data collection and analysis, decision to publish, or preparation of the manuscript.

## Supplementary figures

**Figure S1.**
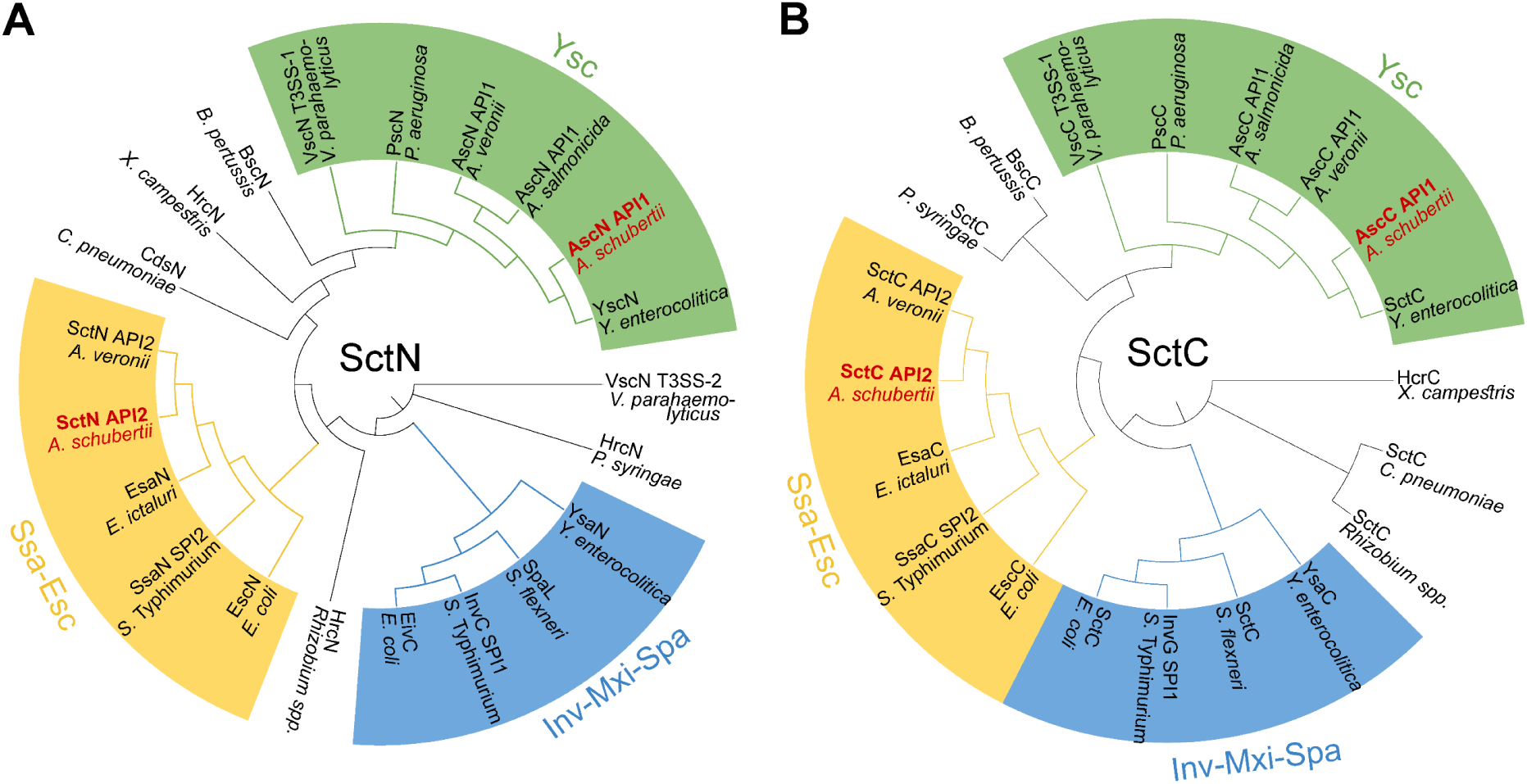
Phylogenetic analysis of core proteins in API1 and API2 injectisomes. To analyze the phylogenetic relationships of SctN and SctC proteins from *Aeromonas* API1 and API2, minimum evolution trees based on *p*-distance were constructed using MEGA11 software. Protein sequences were compared to their orthologs in different bacterial species. The results indicate that AscN and AscC of API1 map within the Ysc family, named after *Yersinia* spp., while SctN and SctC of API2 belong to the Ssa-Esc family, which is characteristic for the SPI2-encoded T3SS injectisome in *Salmonella enterica* serovar Typhimurium.

**Figure S2.**
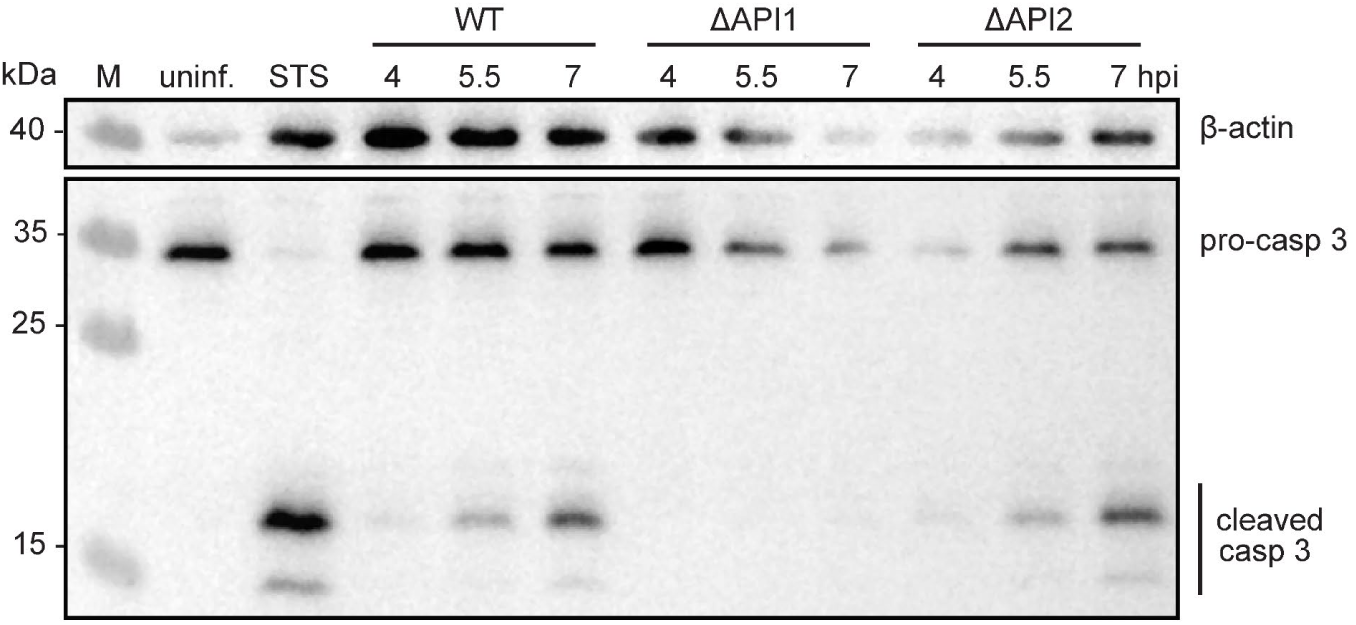
Western blot analysis of caspase-3 activation. HeLa cells were either left uninfected or infected with *A. schubertii* wild-type (WT) or mutant strains lacking the API1 (ΔAPI1) or API2 (ΔAPI2) injectisomes, at MOI of 10:1. One hour post-infection, the extracellular bacteria were eliminated by addition of gentamicin. Whole-cell lysates were prepared at indicated time points, separated by SDS-PAGE and analyzed by immunoblotting using an antibody that detects both the full-length inactive form of caspase-3 (pro-casp 3, 35 kDa), and the cleaved caspase-3 fragment (cleaved casp 3, 17 kDa). β-actin (40 kDa) was used as a loading control. HeLa cells treated with 1 µM staurosporine (STS) served as a positive control. Data are representative of 2 independent experiments.

**Figure S3.**
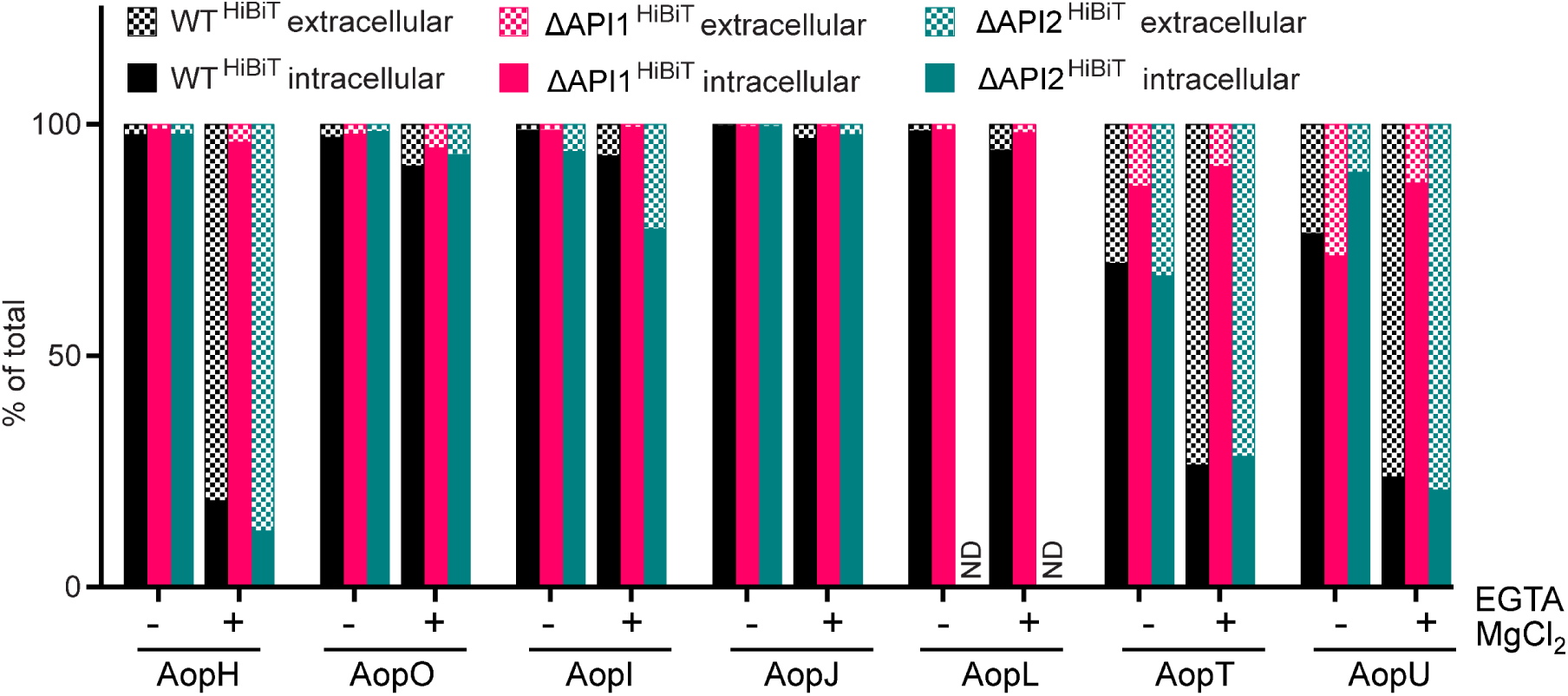
Response of candidate effector proteins to low Ca^2+^/ high Mg^2+^ concentrations. Cultures of WT reporter strains (WT^HiBiT^) and mutant derivatives lacking API1 (ΔAPI1^HiBiT^) or API2 (ΔAPI2^HiBiT^) injectisomes were grown in TSB medium without and with supplementation of 0.5 mM EGTA and 20 mM MgCl_2_. The amount of candidate effector^HiBiT^ in each fraction was expressed as a percentage of the total candidate effector^HiBiT^ present in the culture. Data are representative of 3 independent experiments. ND, not determined.

## Supplementary tables

**Table S1.**
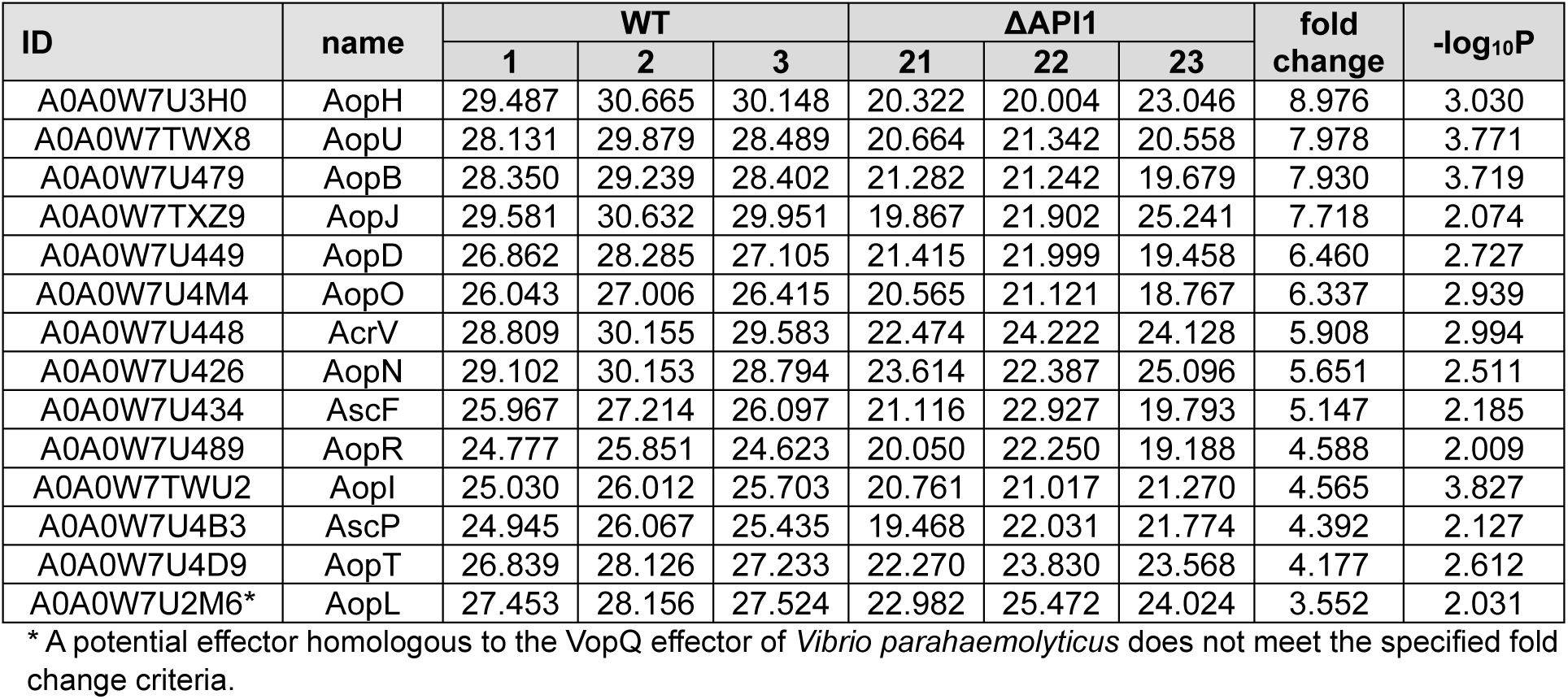
Significant changes in the secretome of WT strain compared to the ΔAPI1 derivative in TSB supplemented with 0.5 mM EGTA and 20 mM MgCl2. This table contains Log2 transformed LFQ proteomic data. Significant changes were defined as |fold change| ≥ 4 and -log10P value ≥ 2, corresponding to P ≤ 0.01.

**Table S2.**
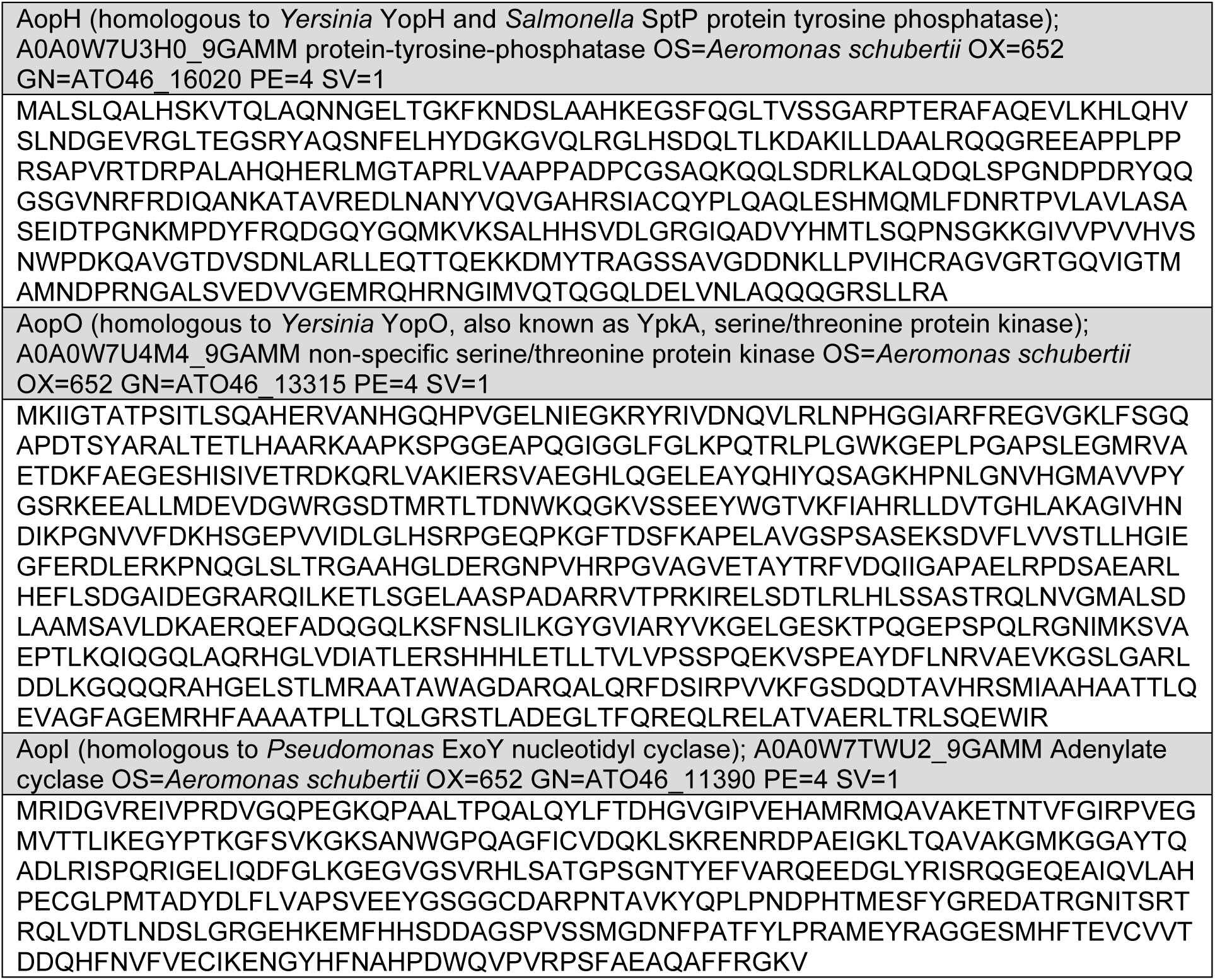
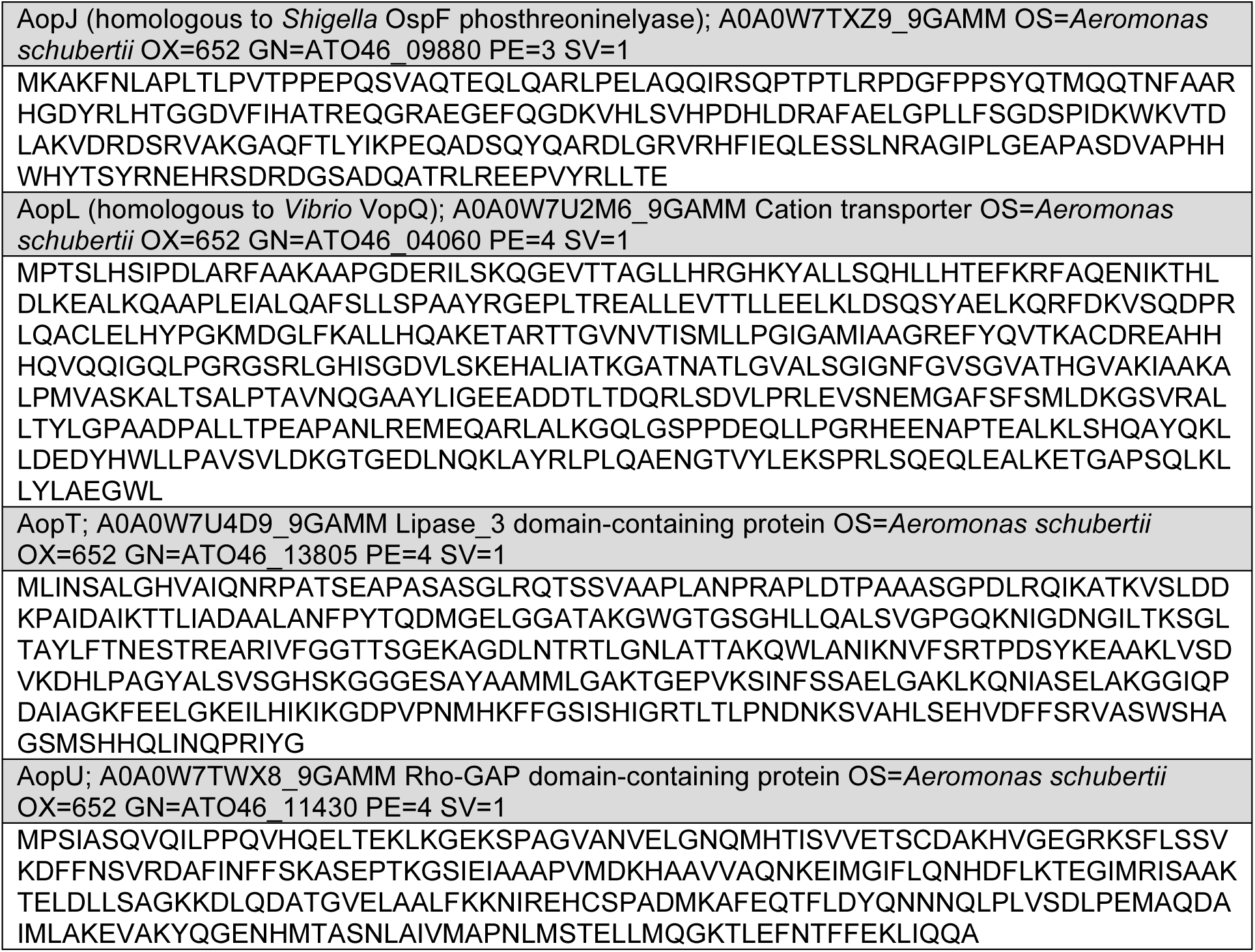
Candidate effectors of API1 injectisome.

**Table S3.**
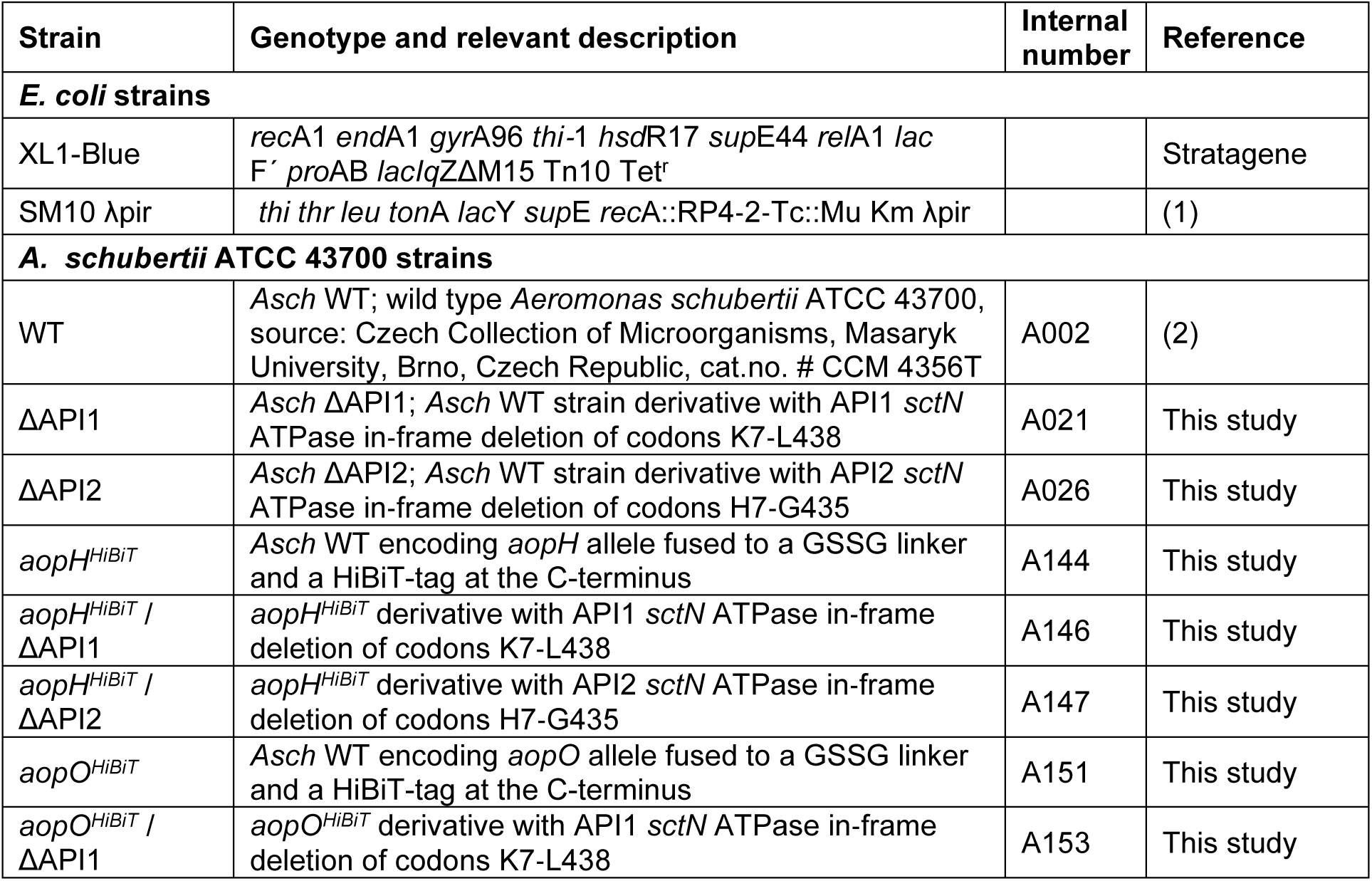
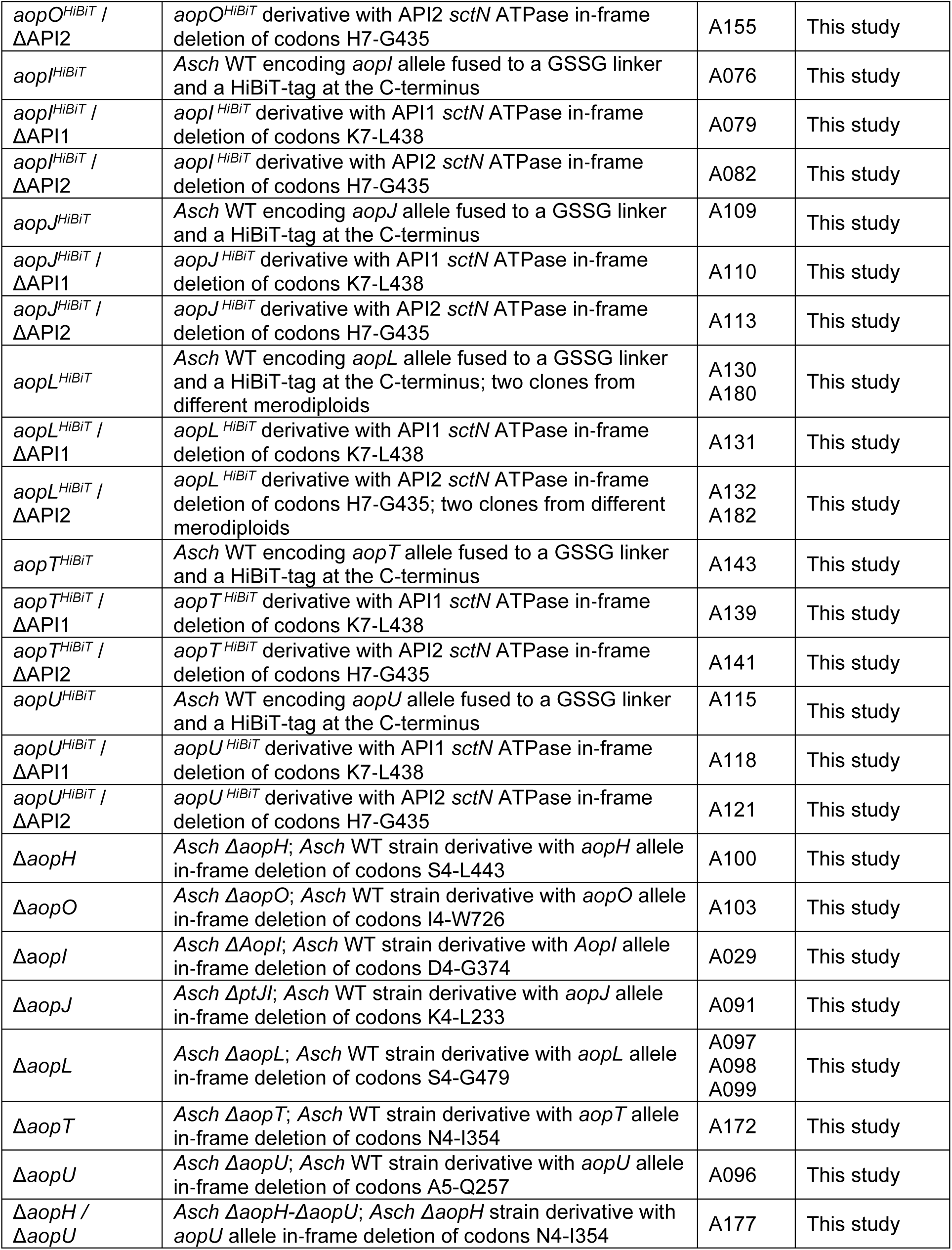
List of bacterial strains used in this study.

**Table S4.**
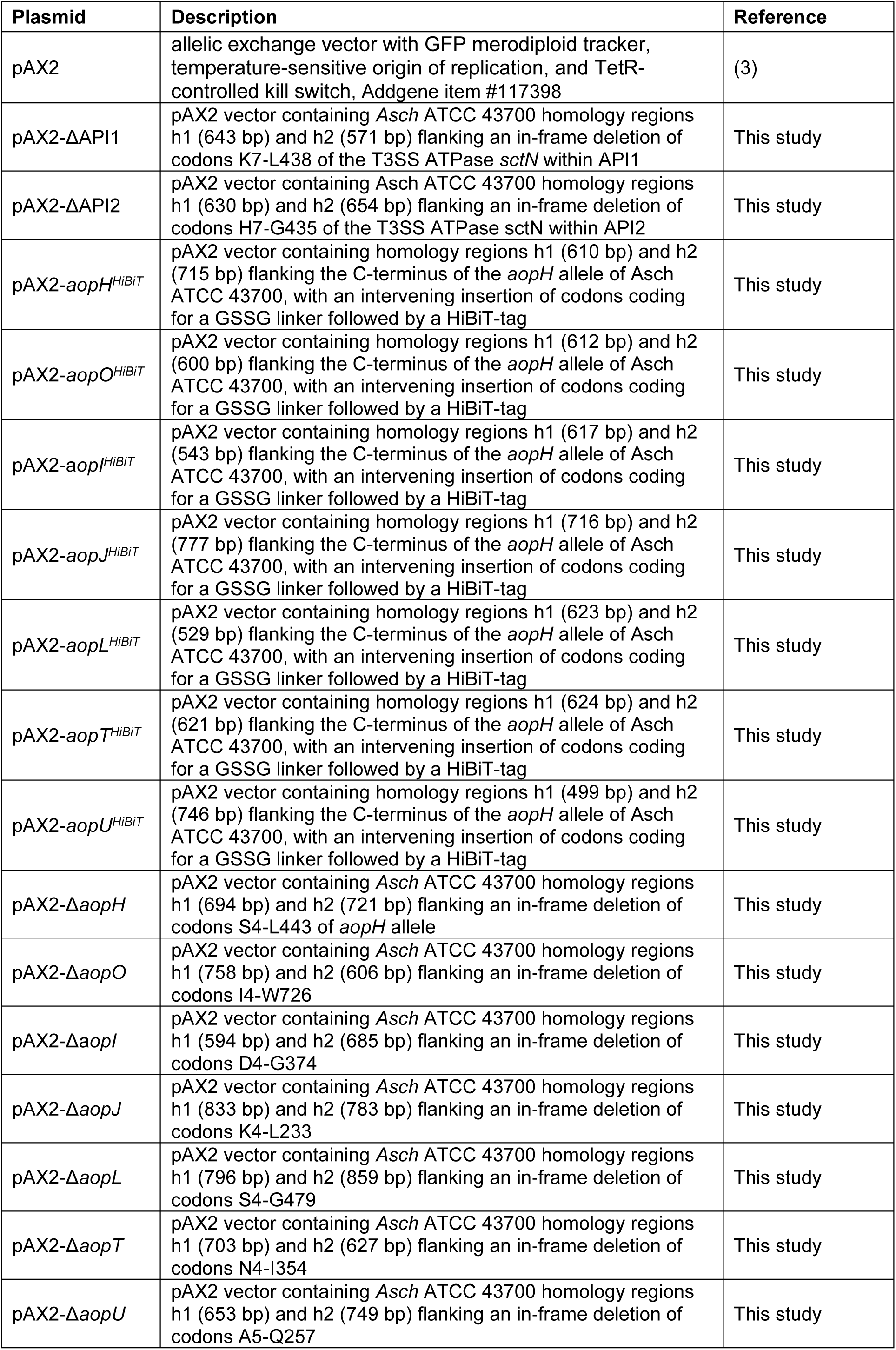
List of plasmids used in this study.

**Table S5.** Accession numbers of SctN and SctC proteins used for phylogenetic analysis.

| Organism | SctN | SctC |
| --- | --- | --- |
| *Aeromonas salmonicida* | ABO92550.1 | CAE83117.1 |
| *Aeromonas schubertii* API1 | WP_050666225.1 | WP_168929881.1 |
| *Aeromonas schubertii* API2 | WP_050667095.1 | WP_019840887.1 |
| *Aeromonas veronii* API1 | MCR4448139.1 | ABP51942.1 |
| *Aeromonas veronii* API2 | WP_021229284.1 | WP_019840887.1 |
| *Bordetella* spp. | WP_003820061.1 | WP_010930813.1 |
| *Edwardsiella ictaluri* | ACR68170.1 | WP_015870340.1 |
| *Escherichia coli* E2348/69 | WP_000622543.1 | AAC38377.1 |
| *Escherichia coli* O157:H7 | WP_000622545.1 | WP_000694687.1 |
| *Chlamydia pneumoniae* | WP_010883345.1 | WP_010883340.1 |
| *Pseudomonas aeruginosa* | WP_003113545.1 | WP_003100753.1 |
| *Pseudomonas syringae* | ABQ88354.1 | WP_103689236.1 |
| *Rhizobium* spp. | ACE93802.1 | WP_065114606.1 |
| *Salmonella enterica* Typhimurium SPI1 | WP_000856766.1 | WP_000848113.1 |
| *Salmonella enterica* Typhimurium SPI2 | WP_000787213.1 | WP_000261284.1 |
| *Shigella flexneri* | WP_000122616.1 | WP_010921673.1 |
| *Vibrio parahaemolyticus* T3SS-1 | WP_005463220.1 | WP_011105906.1 |
| *Xanthomonas campestris* | WP_016851244.1 | WP_016851246.1 |
| *Yersinia enterocolitica* DMO110 | AAB69192.1 | AGM14903.1 |
| *Yersinia enterocolitica* DSM 13030 | WP_010891217.1 | CBY78163.1 |

---

**Supplementary Video S1. Imaging of morphological changes in HeLa cells.**

HeLa cells were either left uninfected or infected with *A. schubertii* wild-type (WT) or mutant strains lacking the API1 (ΔAPI1) or API2 (ΔAPI2) injectisomes, due to the deletion of the respective T3SS ATPases, at MOI of 10:1. One hour post-infection, the extracellular bacteria were eliminated by addition of gentamicin, and morphological changes in HeLa cell were recorded as a time-lapse of 24 hours with frame intervals of 10 minutes. Scale bar, 20 µm.

